# *Agrobacterium*-mediated transient transformation of sorghum leaves for accelerating functional genomics and genome editing studies

**DOI:** 10.1101/779918

**Authors:** Rita Sharma, Yan Liang, Mi Yeon Lee, Venkataramana R. Pidatala, Jenny C. Mortimer, Henrik V. Scheller

**Affiliations:** Joint BioEnergy Institute, Emeryville, CA 94608, USA; Environmental Genomics and Systems Biology Division, Lawrence Berkeley National Laboratory, Berkeley, CA 94720; Crop Genetics and Informatics Group, School of Computational & Integrative Sciences, Jawaharlal Nehru University, New Delhi 110067, India; Department of Plant and Microbial Biology, University of California Berkeley, Berkeley, CA 94720, USA

**Keywords:** *Agrobacterium*, CRISPR, sgRNA, sorghum, transformation, transient

## Abstract

**Objectives:** Sorghum is one of the most recalcitrant species for transformation. Considering the time and effort required for stable transformation in sorghum, establishing a transient system to screen the efficiency and full functionality of vector constructs is highly desirable.

**Results:** Here, we report an *Agrobacterium*-mediated transient transformation assay with intact sorghum leaves using green fluorescent protein as marker. It also provides a good monocot alternative to tobacco and protoplast assays with a direct, native and more reliable system for testing single guide RNA (sgRNA) expression construct efficiency. Given the simplicity and ease of transformation, high reproducibility, and ability to test large constructs, this method can be widely adopted to speed up functional genomic and genome editing studies.

## INTRODUCTION

Sorghum is a gluten-free C4 crop, important as both a human dietary staple and animal feed, but more recently also as a potential feedstock for biofuel production[1]. With high collinearity and synteny with other grass genomes, sorghum also provides an ideal template to serve as model for other grasses[2]. However, realizing the full potential of sorghum as feedstock requires bioengineering efforts aimed at tailoring sorghum biomass for biorefining applications [3, 4]. Indeed, while the sorghum genome sequence was completed a decade ago [2], only a handful of genes have been characterized using transgenic approaches.

A major factor in the lack of progress is the low efficiency and time-consuming nature of stable transformation. Indeed, sorghum is one of the most recalcitrant crops to transformation and regeneration. The first sorghum transgenic plants were generated using particle bombardment in 1993 with only 0.28% transformation rate[5]. Subsequently, Zhao and coworkers[6] reported 2.12% transformation rate using *Agrobacterium*-mediated transformation. Although with recent advancements in technology and optimization of regeneration protocols, several labs have been able to now transform a few limited sorghum cultivars with improved efficiency; reproducibility and consistency still remain major issues[7-9].

When developing engineered plants, due to the time and cost involved, it is highly desirable to test construct functionality in a transient assay. This is particularly true for sorghum. Transient assays in grasses mostly rely on protoplasts[12-14]. However, expression of a gene in protoplasts may not always mimic *in planta* native state and, also experience inconsistent efficiency due to variability in quality of protoplasts and size of vector transformed[15]. Here, we have established a simplified transient assay with *Agrobacterium*, also known as agroinfiltration, for transient transformation of sorghum and demonstrated its application by confirming gene editing in sorghum leaves using GFP as a marker. Using our method, researchers can directly test the *in planta* efficacy of binary constructs that may subsequently be used for stable transformation.

## MAIN TEXT

### METHODS

#### Plasmids and bacterial strains

Binary vectors C282 and C283 were built based on pTKan-p35S-attR1-GW-attR2 backbone vector[16] using Gateway (Invitrogen, CA, U.S.A.) to introduce codons for GFP (C282) or frame-shifted (fs)GFP (C283) for expression under the CaMV 35S promoter. The fsGFP has a 23 bp positive target control (PTC) sequence inserted after the start codon (5’-gcgcttcaaggtgcacatggagg-3’)[21]. C286 contains GFP driven by maize Ubiquitin 1 promoter, described elsewhere[17, 18]. Binary vectors C475 and C476 were built based on pTKan-pNOS-DsRed-pZmUBQ1-attR1-GW-attR2 backbone vector[17]. The C476 cassette (pTKan-pNOS-DsRed-tNOS-pZmUBQ1-CAS9p-pOsU3-PTC_gRNA-p35S-fsGFP) contains a sgRNA (5’-gcgcttcaaggtgcacatgg-3’) targeting the PTC sequence in fsGFP. CAS9p is a plant codon optimized CAS9 from *Streptococcus pyogenes*[19]. The C475 cassette (pTKan-pNOS-DsRed-tNOS-pZmUBQ1-CAS9p-pOsU3-nongRNA-p35S-fsGFP) lacking a sgRNA targeting sequence was used as a negative control. Plasmids are available from the JBEI registry: https://registry.jbei.org

Binary vectors were transformed into *Agrobacterium tumefaciens* strain GV3101 using electroporation, and grown in Luria Bertani (LB) medium containing 100/30/50 μg/mL rifampicin/gentamicin/spectinomycin at 28°C. Similarly, *A. tumefaciens* strain C58C1 containing the P19 suppressor of gene-silencing protein was grown in LB media containing 100/5/50 μg/mL rifampicin/tetracycline/kanamycin.

#### Leaf infiltration

For agroinfiltration, *Agrobacterium* was grown in liquid culture (5 mL, 24 h, 30°C), and cells were pelleted (5000 x*g*, 5 min), and resuspended in infiltration medium containing 50 mM MES, pH 5.6, 2 mM Na_3_PO_4_, 0.5% (w/v) dextrose, 200 μM acetosyringone and 0.01% Silwet L-77 with an OD_600_ of 0.5. The P19 strain was mixed with each of the other strains to ¼ of the final volume. Prior to infiltration, the *Agrobacterium* suspension was incubated without shaking at 30°C for about 2 h. The *Nicotiana benthamiana* plants were grown in a growth chamber under 16/8 h and 26/24°C day/night cycle, and plants of ∼4-weeks-old used for infiltration. *Sorghum bicolor* (L.) Moench inbred line Tx430 plants were grown in a plant growth room under 14/10 h 29/26°C day/night cycle. Plants at the three-leaf stage (3-4 weeks old), were used for co-infiltration (Fig. 1). The fully expanded sorghum leaves were mechanically wounded with a syringe needle (0.8 mm × 40 mm) several times to make the epidermis more conducive to infiltration. No injury was caused to tobacco leaves. The *Agrobacterium* strains suspended in infiltration medium were infiltrated into leaves using a 1 mL syringe without needle. The boundaries of regions infiltrated with *Agrobacterium* were marked with a permanent marker for later visualization.

**Figure 1.**
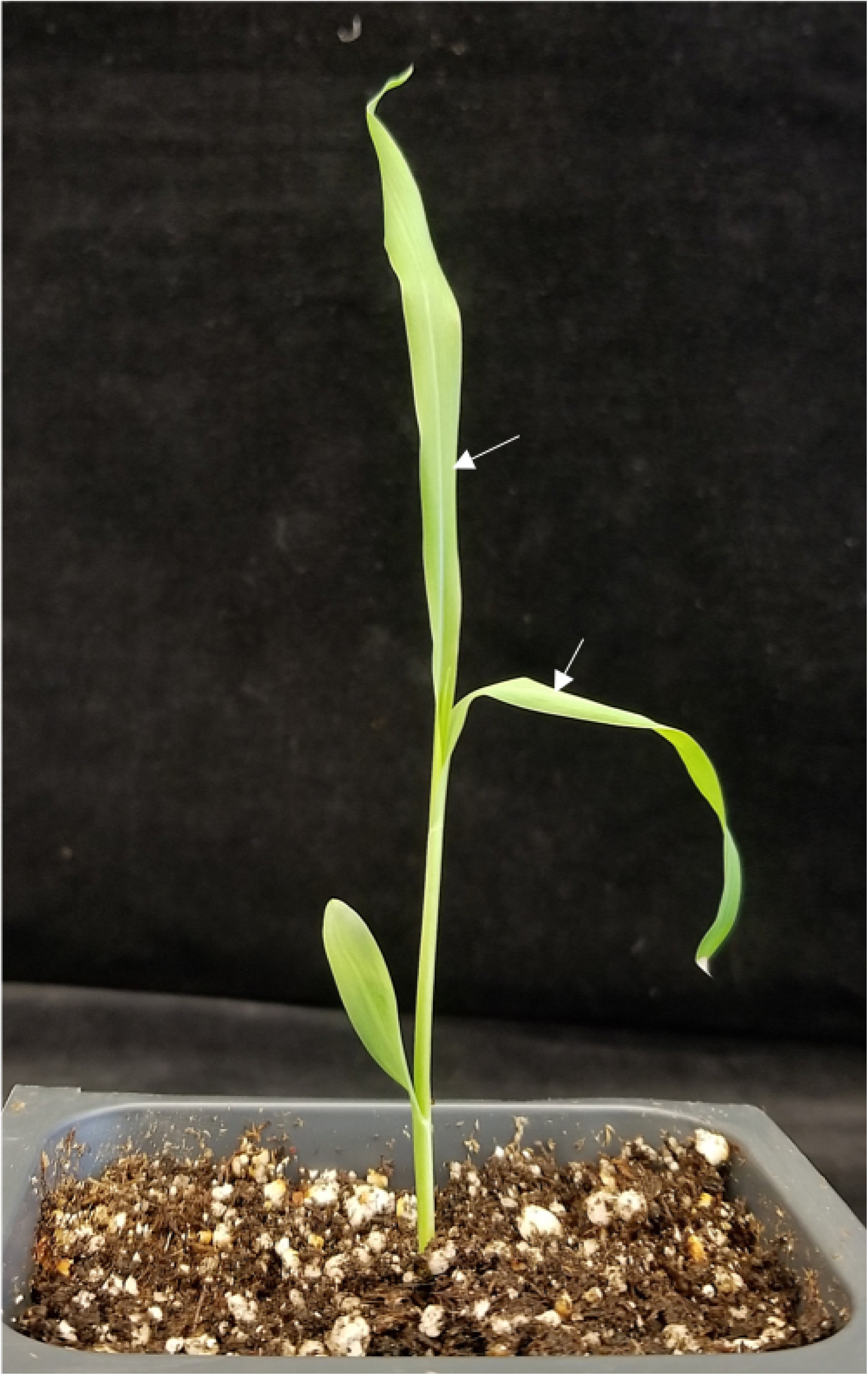
Image of sorghum seedling depicting the stage of sorghum plant required for efficient agroinfiltration. Leaves used for syringe-mediated infiltration on abaxial side are marked by white arrows.

#### Microscopy

About 3-4 days after infiltration (DAI), tobacco and sorghum leaves were detached from the plant and observed under a Leica D4000B fluorescence microscope coupled with a Leica DC500 camera using appropriate filters for GFP and DsRed.

## RESULTS

### Expression of GFP in infiltrated leaves of tobacco and sorghum

We tested binary constructs C282 containing 35S_pro_∷GFP and the modified plasmid C283 with 35S_pro_∷fsGFP (frame-shifted GFP) by agroinfiltration in both tobacco and sorghum leaves. At 3DAI, the GFP signal was examined in detached leaves under a fluorescent microscope. Both sorghum and tobacco leaves infiltrated with C282 showed high and consistent expression of GFP (Fig. 2). However, those infiltrated with C283, containing fsGFP, exhibited no signal. It was noted that the area of detectable GFP expression was much smaller in sorghum as compared to tobacco. This is likely due to the limited infiltration of *Agrobacterium* suspension in sorghum leaves. The signal could be observed up to 7 DAI, after which the signal declined.

**Figure 2.**
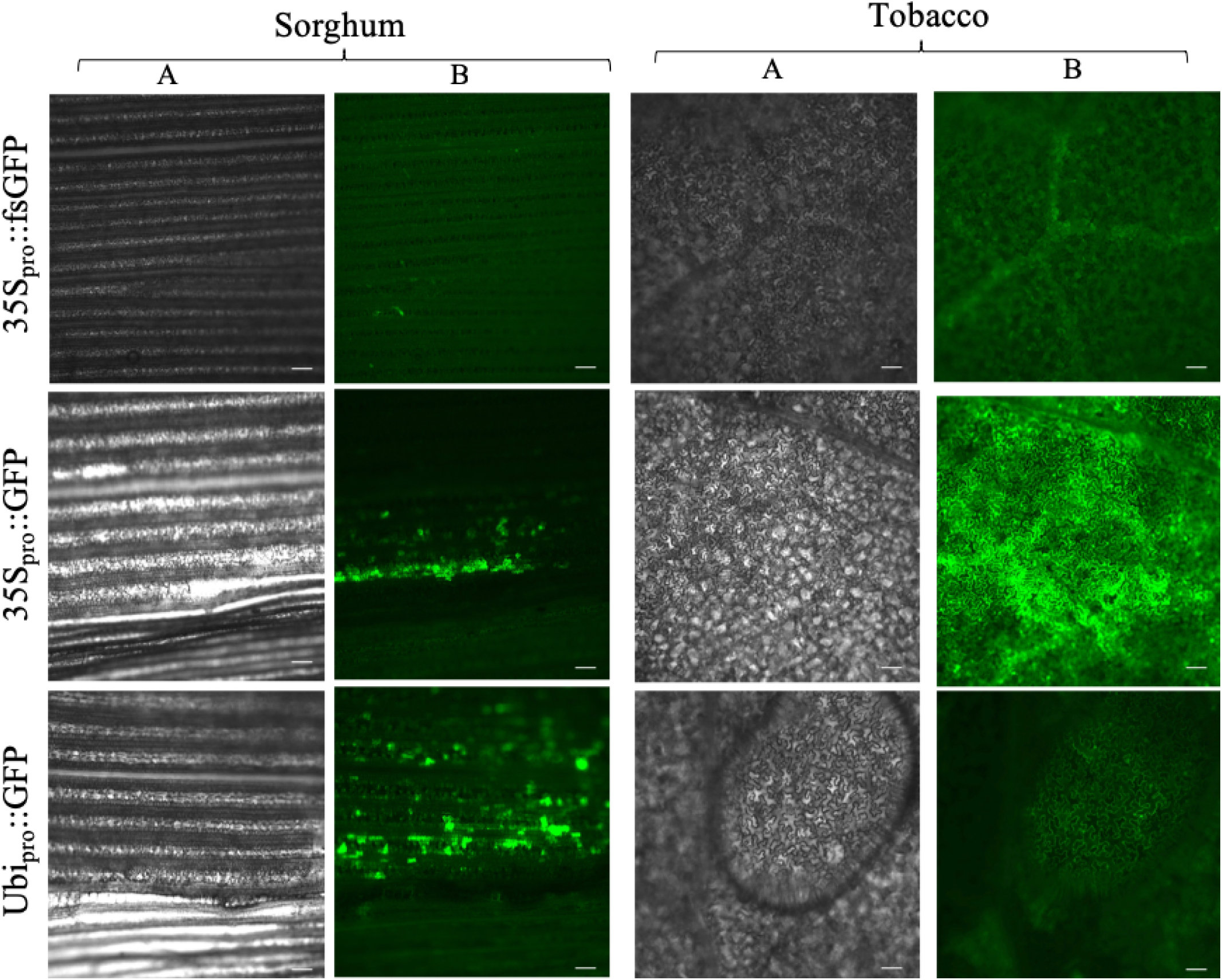
Results of agroinfiltration with *Agrobacterium* suspension in sorghum and tobacco leaves. Column A shows bright field images and column B depicts GFP expression detected using fluorescence microscope. Scale bar: 100 μm.

### Ubiquitin promoter is more effective for sorghum

We compared infiltration of plasmid C282 (35S_pro_∷GFP) with C286 (Ubq_pro_∷GFP) in sorghum. While a higher intensity of GFP signal was observed in tobacco leaves with the 35S promoter compared to sorghum leaves (Fig. 2); GFP expression driven by the maize ubiquitin1 promoter exhibited higher intensity in sorghum leaves.

### Demonstration of gene editing in sorghum leaves using GFP as target gene

To test whether we can use our transient *Agrobacterium*-mediated transformation method to determine sgRNA gene editing efficiency in sorghum, we used the binary vectors, C475 and C476 for agroinfiltration. Tobacco leaves were also infiltrated as a comparison. Both C475 and C476 contained constitutively expressed DsRed under NOS promoter, fsGFP driven by 35S promoter and pUbi-driven CAS9p for CRISPR-mediated genome editing. C476 contained a sgRNA targeting the PTC sequence in fsGFP. As C475 lacked the targeting sgRNA, GFP expression was only expected with C476 vector and only when editing occurs to correct the GFP frame shift.

Following agroinfiltraion, DsRed expression could be detected in both sorghum and tobacco leaves with both the constructs, confirming successful infiltration (Fig. 3). However, GFP expression was observed only in the leaves infiltrated with C476 demonstrating successful editing in the intact leaves of both tobacco and sorghum (Fig. 3).

**Figure 3.**
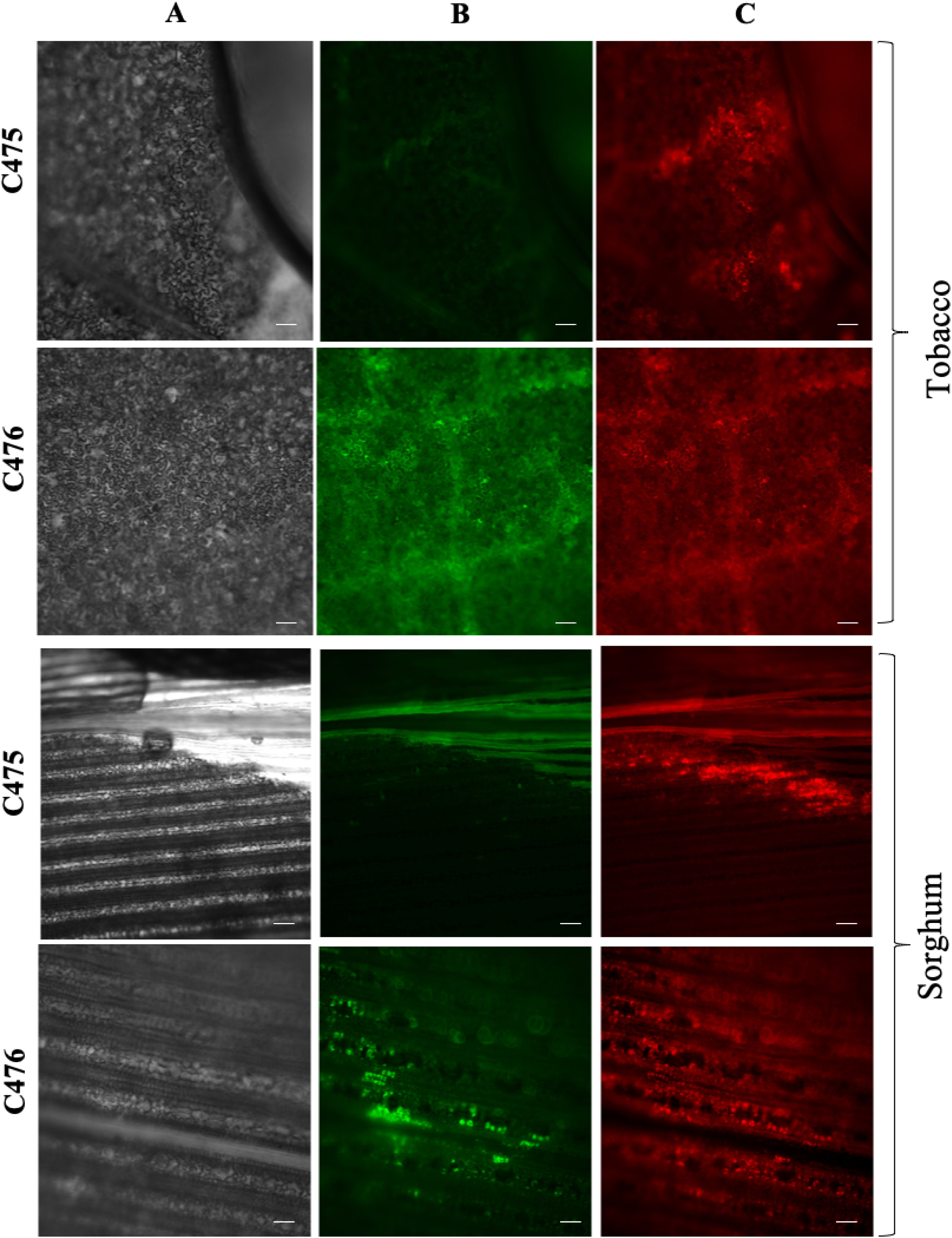
Successful editing of GFP in tobacco and sorghum leaves using agroinfiltration. Column A presents bright field images, whereas, columns B and C present expression of GFP and DsRed, respectively. The C476 vector construct contained sgRNA required for editing, while C475 lacked the sgRNA and serves as negative control. Expression of GFP in leaves transformed with C476 demonstrates successful editing. Scale bar: 100 μm.

## DISCUSSION

Plant transformation is indispensable for elucidating gene function and engineering plant genomes for improved agronomic traits. Several biological, mechanical, chemical and electrical methods of DNA delivery have been developed to facilitate plant transformation over past several decades[22, 23]. Among biological methods, the soil-borne gram-negative bacterium *A. tumefaciens* is no doubt the most popular and widely used vehicle for DNA delivery in plant cells[24]. Although monocots are outside the host range of this bacterium, *Agrobacterium*-mediated transformation is now routinely used for transforming monocot genomes as well, though with lower efficiency[25, 26]. Agroinfiltration is also routinely used in several plant species due to rapidity, versatility and convenience[27-33]. However, success of this method in monocot species is very limited primarily due to extensive epidermal cuticular wax, high silica content, and low volume of intercellular space. These morphological features prevent the infiltration of bacterial cells into grasses *via* the application of simple pressure. Although microprojectile bombardment may be used to introduce expression constructs in cereals, the set-up cost for establishing microprojectile bombardment is high. Moreover, it only targets single cells limiting the scope of screening[34], and often leads to cell damage. Earlier, Andrieu and coworkers[35] reported *Agrobacterium*-mediated transient gene expression and silencing in rice leaves by mechanically wounding leaves followed by direct incubation in *Agrobacterium* suspension. However, we made several attempts to transform sorghum leaves at different stages of development, using their methodology, but could not detect any expression of GFP (data not shown).

Virus-based vectors provide an alternative opportunity for elucidating monocot gene functions. However, instability of the recombinant vector, improper orientation of insert and inconsistency due to inadequate infectivity, inoculation methods, replication/movement of virus in the host, pose serious challenges[36]. Another recent study demonstrated application of nanoparticles in transformation of wheat leaves by combining wounding treatment with syringe infiltration of the nanoparticles[37]. However, the size of plasmid that can be loaded onto nanoparticles is a major constraint due to size exclusion limit of the plant cell wall (∼20 nm).

To overcome these constraints, we attempted syringe infiltration with recombinant *Agrobacterium,* containing vectors for *in planta* GFP expression, at different stages of development in sorghum leaves. As expected, strength of signal in sorghum leaves was higher with the maize ubiquitin promoter as compared to cauliflower mosaic virus 35S promoter, which is reported to perform better in dicots[38]. In our system, although infiltration medium could enter the mature leaves, GFP expression was only detected in the infiltrated younger leaves of 3-4-week-old plants. The expression of GFP seemed to localize to where bacteria were initially infiltrated through mechanical pressure. We did not observe a spread of signal in the adjacent areas, unlike that reported by Andrieu and coworkers[35] for siRNAs in rice. This observation indicated that although bacteria could enter sorghum leaf cells through the wounded regions, they could not passively diffuse to other cells without mechanical pressure in sorghum leaves. We also tried dipping the leaf in *Agrobacterium* suspension after clipping the leaf from the top, as well as wounding by needle, however *Agrobacterium* could not detectably enter the sorghum leaves without applied mechanical pressure.

Further, we demonstrated the application of our method to test the efficiency of sgRNA in genome editing constructs. CRISPR-associated Cas9 is a powerful genome editing tool for engineering plants[39]. Although the design of sgRNAs and preparation of constructs is straightforward, the accuracy and efficiency of the method relies on the choice of sgRNAs[40]. Several *in silico* prediction tools are available to predict the efficiency of sgRNAs based on the sequence features. However, predicted sgRNAs often have vastly different editing efficiencies in planta[18]. Protoplasts have been commonly used to test sgRNA efficiency. However, obtaining high quality protoplasts for genome editing needs extensive standardization, especially for plants such as sorghum. Secondly, additional cloning steps have to be performed to obtain a smaller vector for protoplast transformation. Thirdly and most importantly, the efficiency predicted in protoplasts may not correlate with the efficiency observed in intact plant tissue[40]. Therefore, screening of sgRNAs to achieve high accuracy and efficiency remains a challenge. We adopted our *Agrobacterium*-mediated transient transformation strategy to test sgRNA-mediated editing efficiency in sorghum leaves. The editing was observed in the transformed tissue within three days after infiltration, thereby providing a reliable assay for testing sgRNAs under native conditions.

We used GFP as a reporter in our study as it allows direct visualization in living tissues without being invasive or destructive and does not need any substrate. Gao and workers[41] demonstrated successful use of GFP as marker for stable transformation in sorghum, avoiding use of antibiotics or herbicides. This strategy can be easily applied in our system to quickly assess the full functionality of the vector constructs. For sgRNAs targeting endogenous genes, efficacy can be tested using RT-PCR or sequencing.

Overall, our study demonstrated that *in planta Agrobacterium*-mediated transient expression of transgenes is achievable in sorghum leaves. High reproducibility, simplicity, rapidity and feasibility to transform large constructs, which can directly be used for stable transformation, are the key advantages of our method. Though this method can be used for subcellular localization studies and physiological assays, the ability to test sgRNA targeting efficiency should be of particular interest.

## LIMITATIONS

1. The efficiency of agroinfiltration is much less compared to that observed in tobacco plants and therefore infiltration of more plants may be necessary if significant amount of materials are required for downstream analysis.
2. Since we were targeting a transgene in our editing assays, editing of an endogenous sorghum gene and confirmation of successful editing by sequencing would be an important step to confirm wide applicability of this method.

## DECLARATIONS

### Ethics approval and consent to participate

Not applicable

### Consent for publication

Not applicable

### Availability of data and material

The datasets used and/or analyzed during the current study are available from the corresponding author on reasonable request.

### Author’s Contributions

RS, JCM and HVS conceptualized the study and designed the experiments. RS, YL, VRP and MYL performed the experiments and analyzed the data. RS drafted the manuscript. All authors read and approved the final manuscript.

## Acknowledgements

Not applicable

